# Characterization of novel role for Rab27B in autophagy regulation in colorectal cancer

**DOI:** 10.1101/2024.03.22.586334

**Authors:** Sahida Afroz, Ranjan Preet, Vikalp Vishwakarma, Andrew E. Evans, Dan A. Dixon

**Author notes:** Correspondence: Dan A. Dixon, Department of Molecular Biosciences, University of Kansas, Lawrence, KS, 66045.

## Abstract

**Introduction:** Autophagy is a dynamic, multi-step process that cells use to degrade damaged, abnormal, and potentially harmful cellular substances. While autophagy is maintained at a basal level in all cells, it is activated at a higher level in many cancer cells and promotes tumor growth, anti-tumor immune response, and resistance to cancer therapy. As a result, autophagy is increasingly being recognized to have an important role in cancer progression and emerging as a potential target for cancer therapy. We recently discovered that small GTPase Rab27B, a known regulator of vesicle trafficking and exosome secretion, is also involved in the autophagy process.

**Methods:** Rab27B was knocked out using CRISPR/Cas9 in CRC cell line HCT116. Western blotting, Immunofluorescence, MTT assay, spheroid formation assay, soft agar assay and xenograft studies were performed to analyze the effects of Rab27B deletion on CRC cells.

**Results:** CRISPR/Cas9 deletion or siRNA knockdown of Rab27B in colorectal cancer cells (CRC) showed an abnormal accumulation of autophagy vesicles. Additionally, we observed a significant increase in the autophagy markers LC3-II and p62 by immunocytochemistry and western blot analysis, suggesting a defect in the autophagy flux process. Lysotracker and mCherry-EGFP-LC3 fusion construct indicate an impairment in autophagosome and lysosome fusion when Rab27B is silenced. This defect was rescued by full-length and constitutively active GTP mutant of Rab27B. As autophagy has been shown to have a pro-survival role in tumor growth and stress response, we hypothesized that the observed defects in autophagy flux resulting from Rab27B loss would cause reduced stress response and tumor growth. Indeed, Rab27B knockout reduced cell viability in response to starvation and a 94% reduction in soft agar colony formation. Rab27B deletion also prevented spheroid formation *in vitro.* Finally, to analyze the effect of Rab27B deletion in tumor formation *in vivo*, we performed a xenograft study with wildtype and Rab27B knockout CRC cells, resulting in a dramatic loss of tumor growth (*p*<0.0001) in the KO cells.

**Conclusions:** Together, our results demonstrate a new role of Rab27B in the autophagy trafficking process in CRC. Future studies will focus on investigating the mechanism of how Rab27B functions in the autophagy pathway and whether Rab27B can be targeted as a potential therapeutic strategy for CRC.

## INTRODUCTION

Approximately 4.6% of men (1 in 22) and 4.2% of women (1 in 24) will be diagnosed with colorectal cancer (CRC) in their lifetime, making it the second leading cause of cancer-related death in the US [1]. Although CRC death rates have decreased since 1975 due to screening, new treatments, and changes in risk factors, there will still be an estimated 153,020 new cases of colorectal cancer and 52,550 deaths in 2023 [2]. Cancer cells consistently deal with various stress and challenges like nutrition deprivation, hypoxia, immune response, anti-cancer treatment, etc. They alter different cellular pathways to deal with these stresses and ensure survival and growth. One such pathway is autophagy, which is increasingly recognized to have an essential role role in cancer initiation, maintenance, and resistance to cancer therapy [3–10]. Autophagy is kept at the basal level in all cells, which helps maintain cellular homeostasis and stress responses like energy supply during starvation, destruction of pathogens, etc [11]. But in many cancers, autophagy is activated at a higher level and correlates with poor prognosis basal level in all cells and helps maintain cellular homeostasis and stress responses like energy supply during starvation, destruction of pathogens, etc [11]. But in many cancers, autophagy is activated at a higher level and correlates with poor prognosis [12–14]. In particular, mutations in *KRAS*, an oncogene frequently mutated (30-50%) in colorectal tumors, are associated with higher levels of autophagy [15, 16]. This higher level of autophagy has been shown to promote tumorigenesis by promoting cell survival against various stress by facilitating cancer cell growth, survival in energy crisis, hypoxia, and chemotherapy [17–19]. Autophagy also provides hypoxia, anti-tumor immune response against NK (natural killer) cells, and resistance to cancer therapy [20–23]. Reports show that inhibition of autophagy by autophagy inhibitors re-sensitized resistant cancer cells to chemotherapy and radiation and induced a massive infiltration of NK cells into melanoma tumors [24–26]. These findings led to autophagy being increasingly recognized as a potential target for cancer therapy. So far, the only approved drug targeting autophagy in cancer is hydroxychloroquine (HCQ) [27, 28]. HCQ has been shown to disrupt the autophagic process by blocking the fusion of autophagosomes and lysosomes [29].[27]z. But HCQ has significant limitations, including impairment of necessary lysosomal function, a high concentration required for activity *in vivo*, and a long half-life contributing to side effects. That gives rise to the need for the discovery of new autophagy-targeting drugs.

Given that autophagy is increased in cancer due to stress response to promote cell survival, the vesicle trafficking process related to autophagy must also be regulated to accommodate this increased demand. The Rab proteins are small GTPases that belong to the RAS-like GTPase superfamily and regulate different cellular vesicle trafficking processes [30, 31]. Growing evidence shows various Rab proteins to be involved in the different stages of autophagy [32–34]. Rab1 contributes to the initiation of autophagy, Rab9 modulates the fusion of autophagosomes with late endosomes, Rab7 is implicated in the fusion of autophagosomes with lysosomes, whereas Rab11, Rab30, and Rab37 are essential for starvation-induced autophagy [33, 35–39]. Rab27B is another member of the Rab family that plays a critical role in the endosomal trafficking, exosome secretion, and delivery of secretory granules [40, 41]. Overexpression of Rab27B is seen and associated with poor prognosis and metastasis in various cancers, including colorectal cancer [42–46]. Increased expression of Rab27B can promote proliferation, invasiveness of cells in culture, and invasive tumor growth in xenograft mouse models in breast, pancreatic, colorectal, and lung cancer [47–51]. Recent research suggests Rab27B may also play a role in autophagy. Rab27B localized to autophagosomes under stress conditions, Rab27B KD led to defects in autophagy flux and accumulation of autophagosomes in neuroblastoma cells, and Rab27A/B double KO mice showed an increase in autophagosomes and lysosomes in the lacrymal gland [52–54].

Given that Rab27B overexpression correlates with poor prognosis in CRC and the presence of Rab27B in the autophagosome-lysosome pathway, with growing evidence suggesting high autophagy flux as pro-tumorigenic in CRC indicates Rab27B might have a significant role in CRC tumorigenesis by regulating autophagy process than just promoting exosome secretion. However, Rab27B’s role in the autophagy process in cancer is not studied. Here, we show that depletion of Rab27B in CRC cell lines blocks autophagy flux and results in an accumulation of autophagosomes. Our results indicate that this defect in autophagy flux is caused by a block in the fusion of autophagosomes and lysosomes, indicating that Rab27B plays a role in the fusion step of the autophagic pathway. Additionally, we show that Rab27B deletion inhibits CRC cell growth *in vitro* in 3D culture and *in vivo* xenograft tumor growth, underscoring Rab27B as a potential therapeutic target in CRC.

## MATERIALS AND METHODS

### Immunohistochemistry

#### Tissue microarray and immunohistochemical analysis

Human Formalin-fixed and paraffin-embedded tissue array (CO1801a) containing 90 cases of colon adenocarcinoma with matched 90 adjacent normal colon tissue was purchased from US Biomax, Inc. IHC analysis was performed to detect the expression of Rab27B protein in the microarray samples. The slides were deparaffinized in 2 baths of xylene for 20 min each and rehydrated in graded incubation of 100%, 95%, 80%, and 60% ethanol for 5 min each. For antigen retrieval, first 0.01M sodium citrate buffer was heated to boiling, and then the slides were immersed in the buffer and heated at 95°C for 15 min. The slides were allowed to cool in the buffer for 35 min, followed by 1X TBS washes and incubation in 3% hydrogen peroxide for endogenous peroxidase quenching. The slides were again washed in 1X TBS, blocked in 2.5% BSA in 0.2% Tween 20 for 1h, and incubated at room temperature for 1.5h in primary antibody solution (Rabbit anti-human Rab27B polyclonal antibody, Cat-13412-1-AP1, 1:200 dilution). The slides were then rinsed 3 times with 1X TBS for 5 min each, followed by incubation in secondary antibody (SignalStain Boost IHC Detection Reagent HRP Rabbit #8114)) at room temperature for 30 min. Following TBS rinsing, the peroxidase reaction was visualized by incubation with DAB solution (1 ml DAB Buffered Substrate + 1 drop DAB chromogen, Impact DAB peroxidase substrate kit; Vector-cat# SK-4105). The slides were then counterstained with hematoxylin, dehydrated, and mounted with cytoseal XLT mounting medium (Thermo Scientific cat # 8312-4).

The immunostained slides were evaluated in a blinded manner without prior knowledge of the pathological parameters of the patients. Rab27B expression was monitored in at least 3 randomly selected fields in each section. The immunoreactivity of Rab27B was scored using a semi-quantitative immunoreactive scoring (IRS) system. The staining intensity was categorized into-high staining (score of 3), moderate staining (score of 2), weak staining (score of 1), and negative staining (score of 0). The score for the percentage of positive cells was categorized into->80 % positive cells (score of 4), 51-80% positive cells (score of 3), 10-50% positive cells( score of 2), <10% positive cells (score of 1), and no positive cells (score of 0). The immunoreactivity score (IRS) was calculated by multiplying the staining intensity score and the percentage of positive cells score.

### Cell culture

Colorectal cancer cells (HCT116, HCA7, HT29, SW480, Moser, DLD1, RIE, and Caco2) were purchased from American Type Culture Collection (ATCC) and maintained in Dulbecco’s Modified Eagle’s Medium (DMEM; Corning-USA) supplemented with 10% FBS (R&D systems) with 100 units per ml Penicillin-Streptomycin (Sigma) at 37°C with 5% CO2. For Rab27B siRNA knockdown, HCT116, SW480, HCA7, and DLD1 cells were transfected with predesigned siRNA against RAB27B (Ref# 134127511; IDT, Coralville, IA) or siRNA Control (Cat# AM4635; Ambion, Austin, TX) using the SiQuest (Mirus, *Trans*IT-siQUEST^®^) using previously described procedure. Cells were transfected for 48h and harvested for western blot and imaging.

### Antibodies and Reagents

The primary antibodies used included anti-LC3A/B (D3U4C, Cat # 12741) (1:1000), anti-LC3B (E5Q2K, Cat # 83506) (1:1000), anti-Rab27A (D7Z9Q, Cat # 69295) at (1:1000), anti-Rab27B (E4V30, Cat # 17572), and α-Tubulin Mouse mAb (DM1A, Cat#3873) all purchased from Cell Signaling Technology. The anti-Rab27B (Cat # 13412-1-AP) was purchased from Proteintech Group, Inc and used at (1:1000). Anti-β-Actin (C4, Cat # 0869100-CF) was purchased from MP Biomedicals and used at (1:5000). The anti-Nucleoporin (p62, Cat # 610497) was purchased from BD Biosciences and is used at (1:1000). Secondary antibodies for western blot-IRDye 800CW Donkey anti-Mouse IgG (Cat # 926-32212) and IRDye 680RD Donkey anti-Rabbit IgG (Cat # 926-68073) were purchased from LI-COR.

### Cell Lines, Expression Vectors and Transfections

Rab27B was knocked out in the human colon cancer cell line HCT116 using Rab27B CRISPR/Cas9 KO plasmid (sc-403498; Santa Cruz Biotechnology), followed by insertion of a homology-directed repair (HDR) plasmid (sc-4403498-HDR Santa Cruz Biotechnology) into the double-stranded break. Transfected cells were selected by visible expression of red fluorescent protein (RFP) and puromycin resistance (0.5 ug/ml) (ThermoFisher Scientific) provided by the HDR plasmid. To establish stable knockout clones, individual colonies were isolated by cloning cylinders, expanded in 12 well plates, and screened by western blotting and RT-PCR for Rab27B. Selected knockout cells were maintained in puromycin (0.2 ug/ml) containing media. In vitro experiments were performed using three individual knockout clones, and the xenograft experiment was performed using two knockout clones.

For addback experiments, full-length human Rab27B was synthesized, cloned into pcDNA3.1zeo(+) vector (Gene Universal Inc.), and transfected into Rab27B KO HCT116 cells. Transfected cells were selected by adding Zeocin (500 ug/ml) (?) 48 hrs post-transfection. Stable clones were screened using western blotting and RT PCR.

peGFP-C1 vectors containing full-length Rab27B (GFP-Rab27B, Addgene plasmid #89447) and mutant forms of Rab27B that encoded constitutively active Q78L (GFP-Rab27B Q78L, Addgene plasmid #89448) dominant negative T23N (GFP-Rab27B T23N, Addgene plasmid #89450), dominant negative N133I (GFP-Rab27B N133I, Adddgene plasmid #89449, and geranylgeranyl-binding mutant (GER) (GFP-Rab27B GG, Addgene plasmid #89451) were gifted by Wendy Westbroek (Reference). HCT116 Rab27B knockout cells were transfected with the above plasmids and selected in G418 (500 ug/ml) (Invitrogen) to generate cells stably overexpressing the GFP-Rab fusion proteins. At least three clones of each cell line were used for in vitro experiments to exclude clonal variation.

HCT116 were transfected with pBABE-puro mCherry-EGFP-LC3B plasmid (gift from Jaynta Debnath, Addgene plasmid, #22418) (reference) and selected in puromycin (0.5 um/ml) (ThermoFisher Scientific) to generate cells stably expressing mCherry-EGFP-LC3B fusion protein.

### Fluorescence Microscopy

Immunofluorescence staining was performed following cell signaling immunostaining protocol. Antibodies used were - anti-LC3A/B (D3U4C, Cat # 12741) (1:200) and α-Tubulin Mouse mAb (DM1A, Cat#3873). Briefly, HCT116 and Rab27B KO cells were seeded in 6-well plates with added coverslips and allowed to grow for 48h. Cells were fixed with ice-cold 100% methanol at -20°C for 15 minutes. Fixed cells were rinsed three times in 1X PBS for 5 minutes each and blocked in 3% BSA (in PBS with 0.3% triton X-100) for 1h. Blocking solution was aspirated, and cells were incubated in the primary antibody solution (1:200 in 1X PBS/ 1% BSA/ 0.3% triton X-100) overnight at 4°C. Next day, cells were washed three times in 1X PBS for 5 minutes each and incubated in secondary antibody (?) at RT for 1-2 h. After incubation, cells were washed with PBS and mounted with ProLong™ Diamond Antifade Mountant with DAPI (#p36962, ThermoFisher Scientific). Cells were visualized and imaged with a microscope (?). Quantification of the fluorescence intensity of the images was calculated using ImageJ software. The same procedure was used for imaging Rab27B knocked-down HCT116, SW480, HCA7, and DLD1 cells.

mCherry-EGFP-LC3 expressing HCT116 cells were seeded at 5*10^4^ cells/well density in triplicates in culture slides (#80841, ibidi GmbH, Germany) and incubated under selected conditions for 48h (Control, siRab27B). Cells were treated with 0.5 µm rapamycin for 8h to induce autophagy followed by incubation with 1µg/ml Hoechst at RT for 5 minutes for nuclear staining. Next, cells were fixed with 2% paraformaldehyde for 10 minutes and washed with 1X PBS. Imaging was done using (?). For quantification, spots with both mCherry (red) and EGFP (green) were counted as autophagosomes and spots with only mCherry (red) were counted as autolysosomes.

Lysotracker images were acquired on HCT116 and Rab27B KO cells incubated in required conditions (control and DMEM without glucose) and stained with 70 nM LysoTracker (#L7526, ThermoFisher Scientific) and 1 µg/ml Hoechst. LysoTracker positive punctae were identified and quantified using ImageJ software.

### Transmission Electron Microscopy

HCT116 and Rab27B KO cells were seeded in 6 well plates containing coverslips at 100,000 cells/ well density. After 2 days, the cells were washed and fixed in freshly prepared 4% paraformaldehyde (with 0.5% glutaraldehyde in 0.1 M cacodylate buffer, pH 7.4) at room temperature for 2 h. The fixed samples were sent to Electron Microscopy Research Laboratory (EMRL) at KU Medical Centre for further processing. Images were taken using a JEOL JEM-1400 Transmission Electron Microscope located at KUMC EMRL.

### Western Blot Analysis

For western blot, cells were harvested in SDS lysis buffer (0.5M Tris-HCl pH 6.8, 5% SDS, 10% Glycerol, 5% β-mercaptoethanol, and 0.002% bromophenol blue) and boiled for 5 minutes at 95°C. Total cellular lysates were separated on SDS/PAGE gels and transferred to PVDF membrane (Immobilon^®^-P^SQ^ Transfer membrane, Millipore). The membranes were blocked at RT with 10% milk in TBS with 0.5% tween-20 for 2 hours followed by overnight incubation with primary antibodies at 4°C with gentle agitation.

Scanning was performed using LI-COR Odyssey DLx Imaging System. The membranes were stripped using Restore^TM^ PLUS Western Blot Stripping Buffer and reprobed as and when required. Densitometric quantification of the blots was performed using ImageJ software.

### Messenger RNA Analysis

The total RNA was extracted using the TRIzol reagent (Invitrogen, Carlsbad, CA) and the concentration was measured by NANODROP 2000 (Thermo Fisher Scientific Inc.). cDNA synthesis was performed using 1 µg of total RNA in combination with oligo(dT) and Improm-II reverse transcriptase (Promega, Madison, WI). qPCR analysis was performed as described previously [55] with SYBR green PCR master mix (Applied Biosystems) along with primers to target Rab2B (Rab2B-Fw: CTGCAAATCTGGGATACGGCTG; Rab2B-Re: GCTGCCGGGCATCCTCTAAC), Rab5A (Rab5A-Fw: GAGAGTACCATTGGGGCTGCTT; RAB5A-Re: GGCTGCTTGTGCTCCTCTGTAG), Rab9A (Rab9A-Fw: TTCCCGTGTCGTTTGAGTGC; Rab9A-Re: CCCAACTCCACCATCTCCAAG), Rab11A (Rab11A-Fw: GCAGGGCAAGAGCGATATCGAG; RAB11A-Re: CAGTTCTTTCAGCCATCGCTCTAC), Rab27A (Rab27A-Fw: GGAGGAAGCCATAGCACTCGC; Rab27A-Re: CTTGTCCACACACCGTTCCA) and Rab27B (Rab27B-Fw: GGGACACTGCGGGACAAGA; Rab27B-Re:

TGCTTGCAGTTGGCTCATCC) on 7300 PCR assay system (Applied Biosystems, Foster City, CA). Primers targeting GAPDH (GAPDH-Fw: CCACTCACGGCAAATTCAACGGCA; GAPDH-Re:

TCTCCAGGCGGCACGTCAGATCC) were used as experimental control for normalization. The fold change in mRNA expression levels was normalized using untreated cells.

### Cell Growth Assays

MTT-based cell growth determination kit (Sigma-Aldrich) was used to analyze cell growth over time following previously described protocol [56]. The cells were seeded in 96-well plates at 5000 cells/well density. Next day, 100 ul MTT reagent (0.5 mg/ml) was added to each well and incubated for 6 hours. Next, 100 ul MTT solvent (100 ml Isopropanol, 14 µl HCL, 200 µl 10% NP-40) was added to each well followed by 30 min incubation. Readings of the wells was taken at 570 nm and was repeated for 5 days.

For colony formation assay, cells were seeded at 500 cells/ well in 6-well plates for 10 days. The culture medium was changed every 3 days. After the appearance of visible colony, the cells were washed with PBS, fixed with methanol for 15 min and stained with 0.5% (w/v) crystal violet for 30 min. The colonies were counted manually using a light microscope.

For soft agar colony formation assay, 5000 cells/well were seeded for 3 weeks following a previously described procedure [57]. Cells were washed with PBS and stained with 0.1% crystal violet for 15 minutes. Images were taken on a gel imager with bright filter, and colonies were counted manually.

For spheroid formation assay, cells were seeded at 4 cells/ well density in Costar 96 well plates-Flat bottom, Ultra low adhesion, Polystyrene (Prod ID: PID0250048, Corning) in DMEM/F12 (1:1) 1X (#11330-032, Thermo fisher scientific) with 10mM HEPES, 100 µg/ml EGF and 10 µg/ml FGF. Spheroids were imaged 3 times/week for 3 weeks. Area of the spheroids was calculated using ImageJ software.

### Tumor Growth Xenografts

HCT116 wt and Rab27B KO cells were cultured in DMEM containing 10% FBS and 4.5 g/L glucose and grown for 2 days. On the injection day, cells were trypsinized and counted using KOVA Glasstic Slides. Cell viability was analyzed by trypan blue at the time of counting cells, and plates with more than 98% viability were used for the xenograft study. Next, the cells were centrifuged and resuspended in 50% matrigel in PBS at a density of 1 million cells/100 ul volume. 100 ul (1 million cells) cell suspension (HCT116 wt and two separate Rab27B KO clones) was injected into the flanks of 5-6 weeks old athymic nude mice (The Jackson Laboratory, USA). Four mice per group were used with two tumors/mouse. The mice were weighed three times/week. The injection sites were observed until the tumors were palpable to the touch. Tumor length and width were measured by a digital caliper thrice weekly. Tumor volume was calculated by the formula- [length x (width)^2^/2]. When the tumor volume reached 1500 mm^3^, the mice were sacrificed by isoflurane euthanasia, and tumors were excised, weighed, measured, and photographed next to a ruler. Tumors collected were split into two parts; half of each tissue was snap-frozen (for immunoblotting and qPCR mRNA analysis) and half formalin-fixed and paraffin-embedded for immunohistochemistry (IHC).

## RESULTS

### Expression of Rab27B in CRC

Previous research in our lab identified Rab27B, but not its closely related family member Rab27A, as one of the binding targets of the RNA-binding protein HuR. We found that HuR can bind the 3’ UTR region of Rab27B mRNA, leading to its stabilization and protein overexpression (Unpublished). To investigate Rab27B gene expression status in CRC, we first assessed Rab27B across 275 TCGA CRC tumors and 349 TCGA and GTEx normal tissue using GEPIA (Gene Expression Profiling Interactive Analysis). Rab27B showed increased mRNA expression in tumors compared to normal tissues, whereas Rab27A was highly expressed in normal tissues, and the expression was reduced in tumors (**Figure 1A)**. Next, we examined human colon epithelium samples from 3 individuals and different CRC cell lines for Rab27B and Rab27A expression by qPCR. Consistent with the GEPIA data, Rab27A showed increased expression (4-fold) compared to Rab27B in normal tissue (**Figure 1B**). On the contrary, Rab27B expression was significantly high in CRC cell lines (HCT116, HCA7, SW480, and DLD1) compared to Rab27A (**Figure 1B**). Finally, to analyze Rab27B protein expression, immunohistochemistry (IHC) was performed on a tumor tissue micro-array with 90 cases of colon adenocarcinoma and 90 matched adjacent normal tissue. Most tumor sections (81) were positive for Rab27B in IHC (**Figure 1C**). Rab27B staining was detected primarily in the cytoplasm and the nucleus of carcinoma cells. The tumor sections also showed a significantly higher IRS score than adjacent normal tissue sections (**Figure 1D**). Finally, we analyzed Rab27B protein expression in different CRC cell lines by western blotting. Consistent with qPCR results, Rab27B protein was increased in CRC cell lines HCT116, HCA7, SW480, and DLD1 compared to normal tissue (**Figure 1E**). Together, these results suggest Rab27B is overexpressed in CRC, but Rab27A expression is not altered.

**Figure 1:**
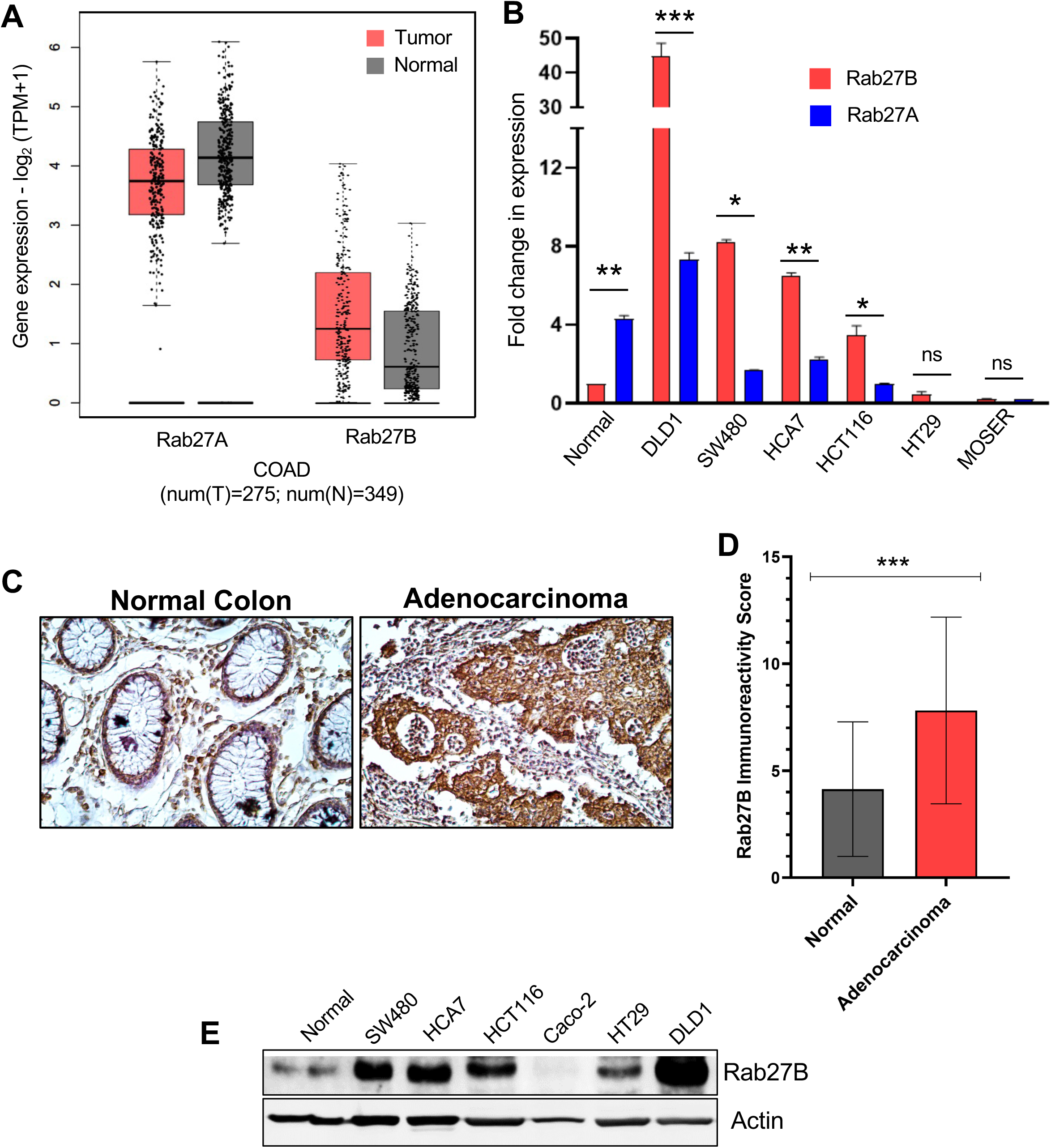
Rab27B is overexpressed in CRC. **(A)** Boxplot derived from gene expression data in GEPIA comparing Rab27A and Rab27B expression in normal tissue and colon adenocarcinoma (COAD) tissue (TCGA database online website GEPIA: http://gepia.cancer-pku.cn/, Normal = 349, Tumor = 275.). **(B)** Analysis of Rab27A and Rab27B expression in normal tissue (n=3) and CRC cell lines (DLD1, SW480, HCA7, HCT116, HT29, and MOSER) by RT-PCR. All data presented are the average ± SEM of three independent experimental repeats. **(C)** Immunohistochemistry images of staged colon adenocarcinoma and adjacent normal tissue from Biomax tissue microarray stained with Rab27B. Best representative images are presented. **(D)** Plot showing immunoreactivity score (IRS) (n=90). **(E)** WB showing the comparison of Rab27B expression in normal tissues vs. CRC cell lines (SW480, HCA7, HCT116, Caco-2, HT29, and DLD1). Actin was used as the loading control. (*, *p* ≤ 0.05; **, *p* ≤ 0.01; ***, *p* ≤ 0.001).

### Effects of Rab27B depletion on CRC cells

To characterize the role of Rab27B in CRC, we performed siRNA-mediated knockdown (KD) of Rab27B in two CRC cell lines, HCT116 and SW480 (**Figure 2A, 2C**), and observed striking phenotypic differences in the KD cells. The KD cells were larger in both cell lines and showed an abnormal accumulation of vesicle-like structures (**Figure 2B, 2D**). Next, we created CRISPR/Cas9 mediated knockout (KO) of Rab27B in HCT116 cells. Knockout was accomplished using the Rab27B CRISPR/Cas9 KO plasmid that expresses three short guide RNA (sgRNA) and the Cas9 endonuclease to create double-stranded breaks. A homology-directed repair (HDR) plasmid was used to insert a gene cassette expressing puromycin resistance and dsRed protein into the double-strand break. Rab27B KO was verified through western blotting for Rab27B protein (**Supplementary Figure 1A**, **Figure 2E**). Consistent with our previous data, the KO cells also showed similar phenotypic changes of increased cell size and accumulation of vesicles (**Figure 2F**). Rab27A expression was not changed in the KO cells (**Supplementary Figure 1B**). Overall, our data showed that Rab27B depletion causes increased cell size and accumulation of vesicles in CRC cells.

**Figure 2:**
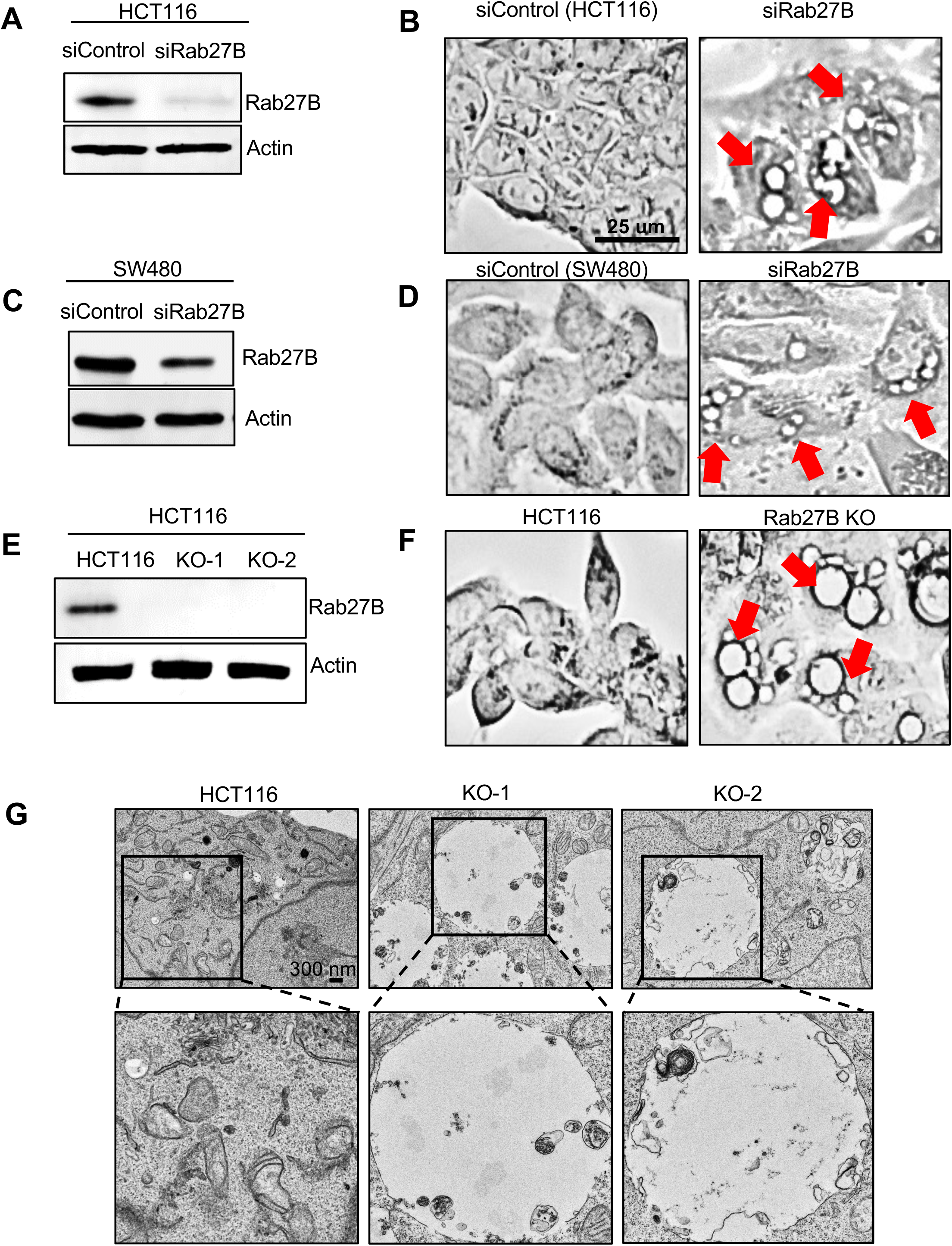
Rab27B suppression results in vesicle formation in CRC cell lines. **(A-D)** Depletion of Rab27B by siRNA-mediated knockdown resulted in the formation of vesicle-like bodies (indicated by red arrows) in HCT116 **(A, B)** and SW480 cells **(C, D)**. **(E)** Rab27B was knocked out by CRISPR/Cas9 in HCT116 cells, resulting in vesicle formation in the KO cells **(F)**. Knockdown and knockout were verified by WB using actin as the loading control. Brightfield images were taken to observe cell morphology. All data presented are representative of four independent experiments. **(G)** Electron microscopy images of HCT116 WT and Rab27B KO cells. Rab27B KO cells showed an accumulation of autophagosome-like vesicles. Images are representatives of two independent experiments.

### Analysis of autophagy defects in Rab27B deleted CRC cells

We used transmission electron microscopy (TEM) to identify and characterize the vesicles observed in Rab27B KO cells. TEM showed the vesicles were larger than 500 nm in area (data not shown), possess double membrane and cytoplasmic contents for degradation, which are characteristics of autophagic vesicles (**Figure 2G**) [58]. To check whether these are autophagic vesicles, we used CYTO-ID Autophagy detection kit, which selectively labels autophagic vesicles at different stages [59]. WT cells treated with rapamycin and chloroquine were used as positive control. Rapamycin induces autophagy, and chloroquine blocks autophagic degradation, which results in an accumulation of autophagic vacuoles in the cell. CYTO-ID staining showed a 2-fold increase in CYTO-ID positive vesicles (green) in KO cells compared to untreated WT cells (**Figure 3A**). We also measured the fluorescence intensity (Absorbance 499/548 nm) of CYTO-ID staining, which showed a significant increase in the KO cells (**Figures 3BC**). From this experiment, we hypothesized that Rab27B depletion results in the accumulation of autophagic vesicles. To test this hypothesis, we looked at the level of the most widely monitored autophagic marker, the LC3 protein [58]. LC3 is a soluble protein distributed ubiquitously in mammalian tissues and cultured cells. During autophagy, the 17 kDa cytosolic form of LC3 (LC3-I) gets cleaved into a 15 kDa (LC3-II) form, conjugates to phosphatidylethanolamine, gets recruited to autophagosomal membranes, and is recycled by the degradation of autophagosomes [60]. Increased levels of the cleaved LC3-II form indicate an increased presence of autophagosomes. First, we wanted to see if there was an increase in LC3 by immunofluorescence staining. LC3-I and LC3-II can be distinguished in immunofluorescence staining by their distribution. A diffused cytosolic distribution indicates LC3-I, whereas intense puncta-like structures indicate autophagosome localized LC3-II [60, 61]. We found that the Rab27B KO cells showed a significant increase in LC3-II punctate structures (**Figure 3D**). We also counted the number of LC3 positive vesicles per cell and saw a significant increase in the KO cells compared to the wt cells (**Figure 3H**). To confirm whether this LC3 increase is consistent across other Rab27B depleted CRC cell lines, we knocked down Rab27B in SW480 and DLD1 cells and saw a similar increase in LC3-positive vesicles in immunofluorescence staining (**Figures 3E and 3I; Supplementary** Figures 2AB). Next, we performed a rescue experiment by transfecting KO cells with a vector containing full-length Rab27B and generated stable addback cell lines. These cell lines showed rescue of the autophagy phenotype in IF (**Figures 3F and 3J**). Interestingly, knocking down Rab27A did not change LC3 level or localization in HCT116, indicating that Rab27A does not play any role in autophagy (**Figures 3G and 3K; Supplementary Figure 3C**). Our findings indicate a dysregulation in the autophagy flux process in the absence of Rab27B and implicate a possible role of Rab27B in the autophagy process in CRC.

**Figure 3:**
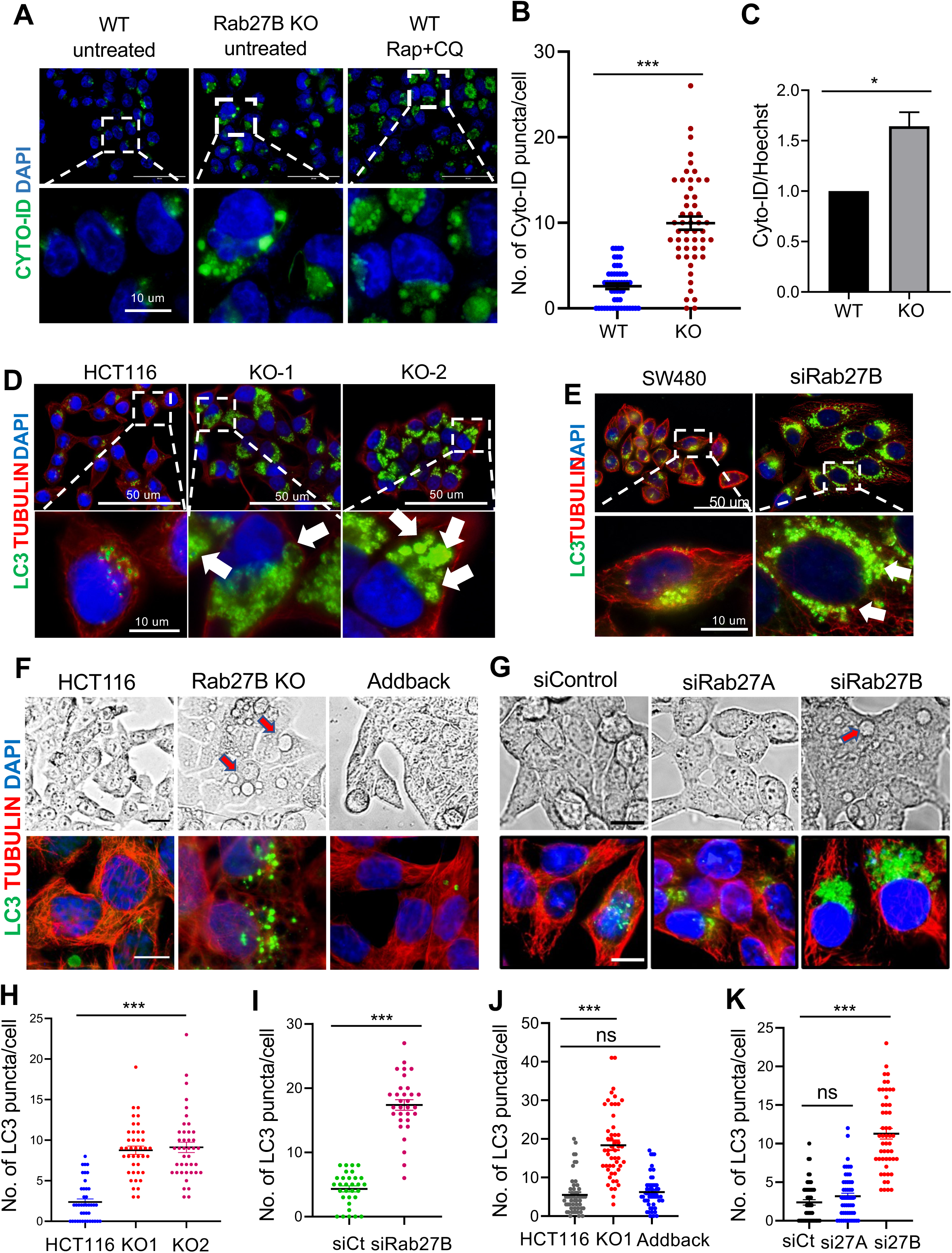
Rab27B KO cells show upregulation of autophagy markers in IF stainings. **(A)** Immunofluorescnece images of HCT116 WT and Rab27B KO stained with CYTO-ID dye to label autophagic vesicles and Dapi to stain the nucleus. WT cells treated with rapamycin (0.5 um) and chloroquine (10 um) overnight were used as positive control. **(B)** Comparison of CYTO-ID positive vesicles/cell in WT and KO cells. **(C)** Absorbance (499/548 nm) for CYTO-ID was measured. **(D)** HCT116 and Rab27B KO cells were immune stained with anti-α-tubulin (red signal) and anti-LC3 (green signal) antibody. Higher magnification images of the area outlined in white show the presence of punctate LC3-II positive structures in Rab27B KO cells (white arrows). Dapi (blue) was used to label the nucleus. **(E)** siRNA-mediated knockdown of Rab27B in CRC cell line SW480 was followed by immunofluorescence staining. Number of LC3 positive vesicles per cell was scored in HCT116 Rab27B KO **(H)** and siRab27B SW480 cells **(I)**. **(F)** Rab27B KO cells were transfected with pcDNA3.1zeo(+) vector containing full-length human Rab27B (Gene Universal Inc.). Transfected cells were selected in media containing Zeocin (500 ug/ml), and stable addback clones were selected and expanded. **(F, J)** Immunofluorescence staining and scoring for LC3 showed rescue of autophagy phenotype in the addback cells. **(G)** Immunofluorescence staining of HCT116 control, siRab27A, and siRab27B cells with anti-α-tubulin (red signal) and anti-LC3 (green signal) antibodies**. (K)** Rab27A KD did not result in autophagic vesicle accumulation, as seen in Rab27B KD cells. Data presented are representative of three independent experiments (*, *p* ≤ 0.05; **, *p* ≤ 0.01; ***, *p* ≤ 0.001).

### Functional implication of Rab27B in the fusion of autophagosomes and lysosomes

Our results indicate that Rab27B silencing leads to an abnormal accumulation of autophagic vesicles in CRC cells. This accumulation can be caused by an increase in autophagy induction or a block in the degradation of autophagosomes. To identify at which step of autophagy Rab27B functions, we first tested whether there was an increase in the induction of autophagy. An increase in autophagy flux can be tested by looking at the LC3-II/LC3-I ratio. A decrease in the ratio indicates an increase in autophagy induction, whereas an increase in the ratio indicates a block in autophagy degradation [58]. First, We performed western blot analysis to compare LC3II/LC3I in HCT116 and Rab27B KO cells (**Figure 4A**). We saw a 3.5 to 4-fold increase in the LC3-II/LC3-I ratio, indicating the latter is occurring (**Figure 4B**). We confirmed this result in Rab27B KD HCT116 and SW480 cells and observed a 2.5-fold increase in LC3-II/LC3-I ratio (**Figures 4C-F**). Adding back Rab27B restored the LC3-II level in the KO cells (**Figure 4G**). Next, to check whether there is any dysregulation in the expression of the genes involved in the autophagic pathway, we conducted RNA-seq analysis on Rab27B KO cells. The heatmap comparing gene expression between wt and KO cells showed no major difference (**Figure 5A**). We also looked at the expression of autophagy-related genes ulk1, ulk2, FIP200, ATG13, ATG14, Beclin1, VPS34, ATG7, ATG3, ATG5, ATG12, STX17, SNAP29, VAMP8, and GABARAP and did not see any significant change in the KO (**Figure 5B**). Given that Rab27B functions in cellular vesicle trafficking, Rab27B KO CRC cells show an accumulation of autophagosomes and an increase in LC3-II/LC3-I ratio, it is probable that Rab27B functions in the autophagy process after the formation of autophagosomes; thus, in the later stages of autophagy. To test this, we looked at the level of p62, a marker of autophagic degradation [62]. p62, also known as sequestosome-1, is a protein that binds to LC3-II and gets degraded by autophagy [63]. An increase in the p62 level indicates a block in the later steps of autophagy, preventing autophagosome degradation [58]. In order to compare the level of p62, we performed immunofluorescence staining in HCT116 wt and Rab27B KO cells for p62 (**Figure 5C**). Rab27B KO cells showed a greater percentage of cells with p62-positive vesicles (60%) compared to control (20%) and a 1.5-fold increase in p62 fluorescence intensity/cell (**Figures 5DE**), indicating a block in the autophagic flux in the KO cells.

**Figure 4:**
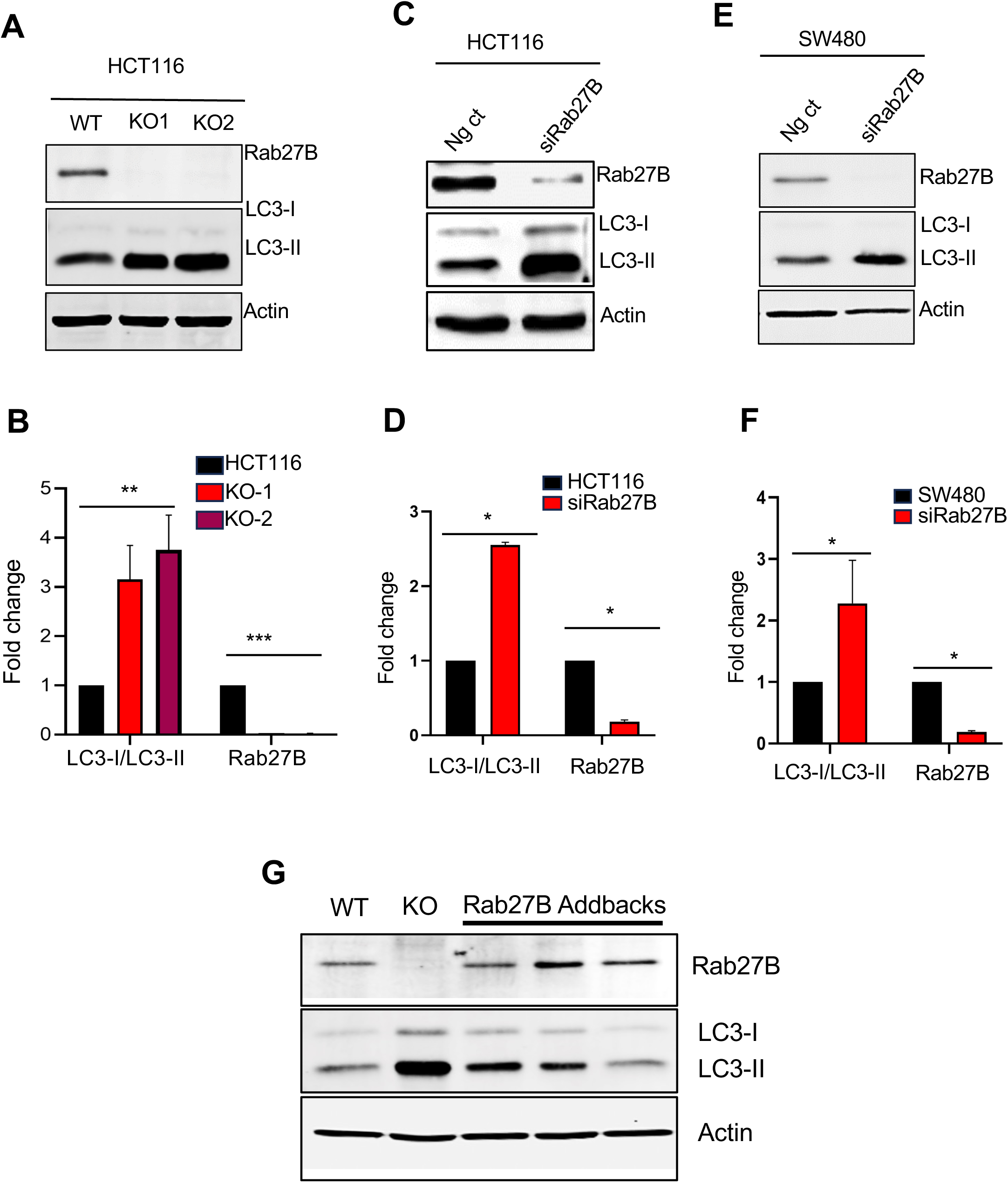
Rab27B depletion increases LC3-II/LC3-I ration. **(A)** HCT116 and Rab27B KO cell lysates were probed for LC3-II by western blot analysis. Rab27B KO cells showed increased LC3-II compared to HCT116. Cell lysates were also probed for Rab27B to verify the absence of Rab27B in the KO cells. Actin was used as the loading control. **(C, E)** WB showing LC3 level in Rab27B KD HCT116 and SW480 cells. **(B, D, F)** Densitometry analysis of LC3-II/LC3-I ratio using ImageJ software. LC3-II/LC3-I was increased in Rab27B depleted cells. **(G)** Western blot of HCT116 wt, Rab27B KO, and Rab27B addback cells shows restoration of LC3-II level in addback cells.

**Figure 5:**
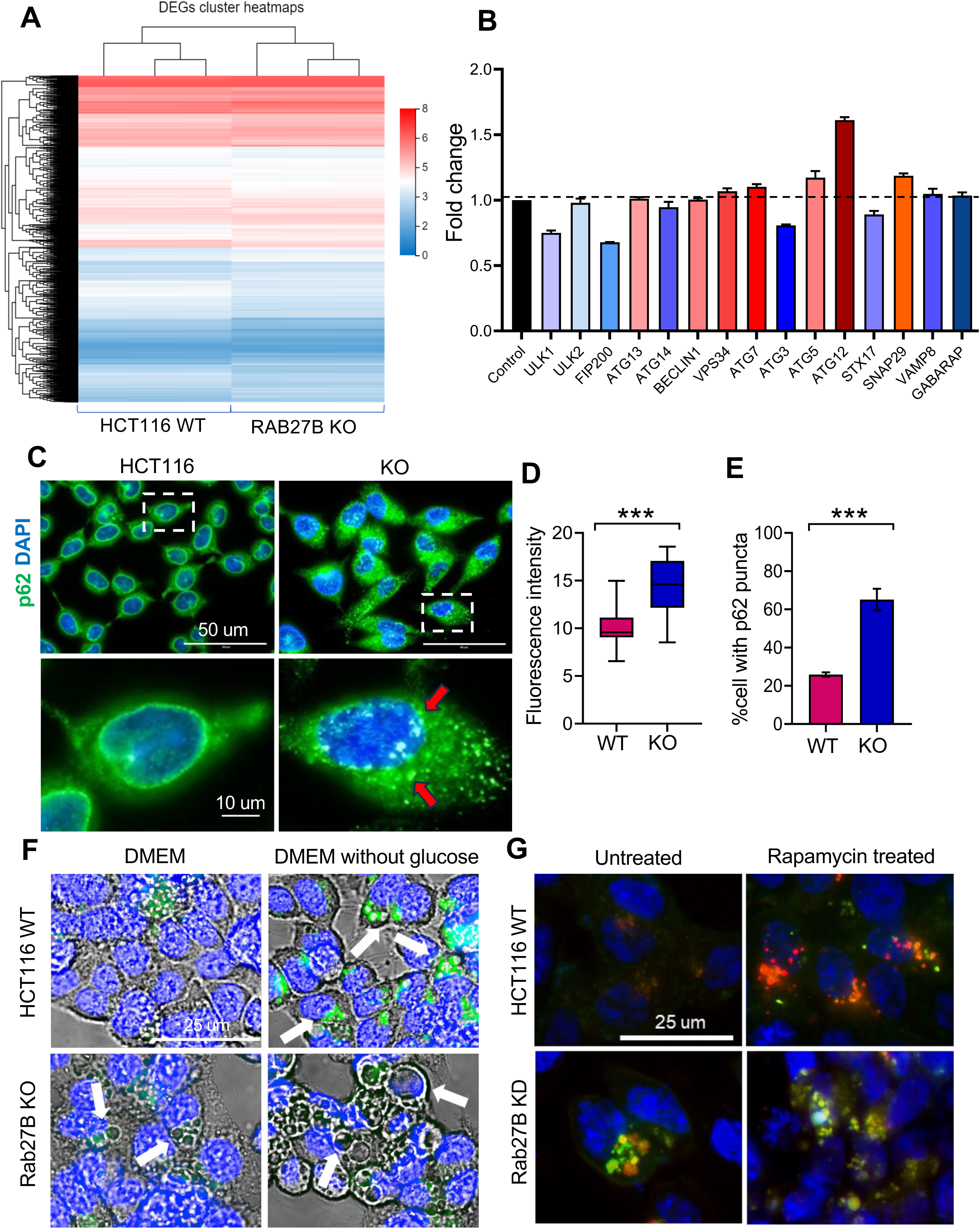
Rab27B deletion impairs autophagy flux. **(A)** Heatmap of RNA seq transcriptome analysis of HCT116 WT and Rab27B KO cells. The scale at the right indicates relative expression levels from high (red) to low (blue). **(B)** Comparison of mRNA expression of different autophagy genes (ULK1, ULK2, FIP200, ATG14, BECLIN1, VPS34, ATG7, ATG3, ATG5, ATG12, STX17, SNAP29, VAMP8, and GABARAP) between WT and KO cells obtained from RNA seq analysis. **(C)** IF staining of HCT116 and Rab27B KO cells with autophagy marker p62 (green) and Dapi (blue). The KO cells exhibited increased oveall p62 fluorescence intensity **(D)** and a higher percentage of cells with p62 positive vesicles **(E)**. **(F)** Lysotracker staining of HCT116 and Rab27B KO cells to label autolysosomes. Dapi was used for nuclear staining. Cells kept in DMEM without glucose (12h) were used as positive controls. Rab27B KO cells showed reduced lysotracker-positive vesicles (indicated by white arrows), suggesting impairment in lysosomal fusion. **(G)** Rab27B was depleted by siRNA-mediated KD in HCT116 cells stably expressing mCherry-EGFP-LC3B. Autophagy flux was monitored by comparing the ratio of mCherry and GFP double-positive vesicles (yellow) and mCherry-positive (red) vesicles. Cells treated with rapamycin (0.5um for 12h) were used as positive controls. Data presented are representative of three independent experiments (*, *p* ≤ 0.05; **, *p* ≤ 0.01; ***, *p* ≤ 0.001).

### Rab27B depletion impairs lysosomal fusion of autophagy flux

We used mCherry-EGFP-LC3 reporter to view free autophagosomes and autophagosomes fused with lysosomes [61, 64, 65]. LC3-II is incorporated into the inner and outer membranes of expanding and mature autophagosomes, which ultimately fuse with lysosomes for degradation. mCherry-EGFP-LC3 labels two stages of autophagic vesicles: autophagosomes that are double-positive for mCherry and EGFP-fluorescence and autolysosomes that, due to pH sensitivity of GFP, are marked solely by mCherry fluorescence (**Supplementary Figure 3A**). An increase in the double-positive mCherry and GFP-fluorescence in the Rab27B KO cells indicates a block at the fusion stage. In contrast, an increase in the mCherry positive fraction only suggests a block in the autolysosome stage. First, we stably expressed mCherry-EGFP-LC3 reporter in HCT116 cells. Untreated cells show cytosolic distribution of double positive LC3, autophagy induction with rapamycin results in the formation of increased autolysosomes (mCherry positive vesicles), and blocking autophagy flux by chloroquine treatment shows increased autophagosomes (EGFP and mCherry double-positive vesicles) (**Supplementary Figure 3B**). Next, we knocked down Rab27B in that cell line. We used rapamycin-treated cells as positive control. Rab27B KD cells showed an increased presence of mCherry-EGFP positive puncta compared to wt cells (**Figure 5G**). Similarly, rapamycin-treated wt cells showed an increased ratio of mCherry positive red puncta, whereas Rab27B KD cells showed an increased percentage of mCherry-EGFP-positive yellow puncta (**Figure 5G**). This increase in the double-positive mCherry and GFP-fluorescence in the Rab27B KO cells again indicates a block at the fusion stage. Next, to investigate whether Rab27B promotes autophagosome degradation by directly trafficking autophagosomes to lysosomes or indirectly by affecting lysosomal biology, we used lysotracker to look at lysosome acidification and distribution. LysoTracker is an acidotropic dye that stains cellular acidic compartments, including lysosomes and autolysosomes [66–68]. Live cells were stained with LysoTracker and Hoechst dye (nuclear stain) for 5 min prior to immediate imaging with an Epifluorescence microscope. Cells starved in DMEM without glucose to induce autophagy were used as positive control. There was an significant increase in lysotracker positive green vesicles in starved wt cells, where as the vesicles in the starved KO cells remained Lysotracker negative (**Figure 5F**). Taken together these results indicate that Rab27B is involved in the autphagosome degradation step in the autophagy process in CRC.

### Requirement of GTPase activity of Rab27B for the autophagic role

Since Rab27B is a GTPase and functions by switching between active GTP and inactive GDP state, we wanted see whether GTPase function of Rab27B is required for the autophagy role. We utilized vectors containing mutant forms of Rab27B as follows-constitutively active GTP mutant Q78L, dominant negative GDP mutant N133I, and T23N, and Geralyn-geralyn mutant GER along with WT form of Rab27B (**Figure 6A**). We expressed these vectors in Rab27B KO HCT116 cells and generated stable cell lines (**Figure 6B**). Brightfield images showed restoration of autophagy phenotype only in the WT and Q78L Rab27B expressing cells (**Figure 6C**). KO cell lines expressing dominant negative Rab27B-T23N and Rab27B-N133I, or geralyngeralynation mutant Rab27B-GER showed accumulation of autophagic vesicles (**Figure 6C**). This result indicates that GTPase activity of Rab27B is necessary for the autophagy function. We also compared the proliferation rate of the mutant Rab27B expressing cell lines by MTT assay. Interestingly, constitutively active GTP mutant Q78L cells showed significantly increased proliferation compared to WT or KO cells (**Figure 6D**), whereas cells expressing Rab27B-T23N or Rab27B-GER did not change in growth (**Figure 6E**).

**Figure 6:**
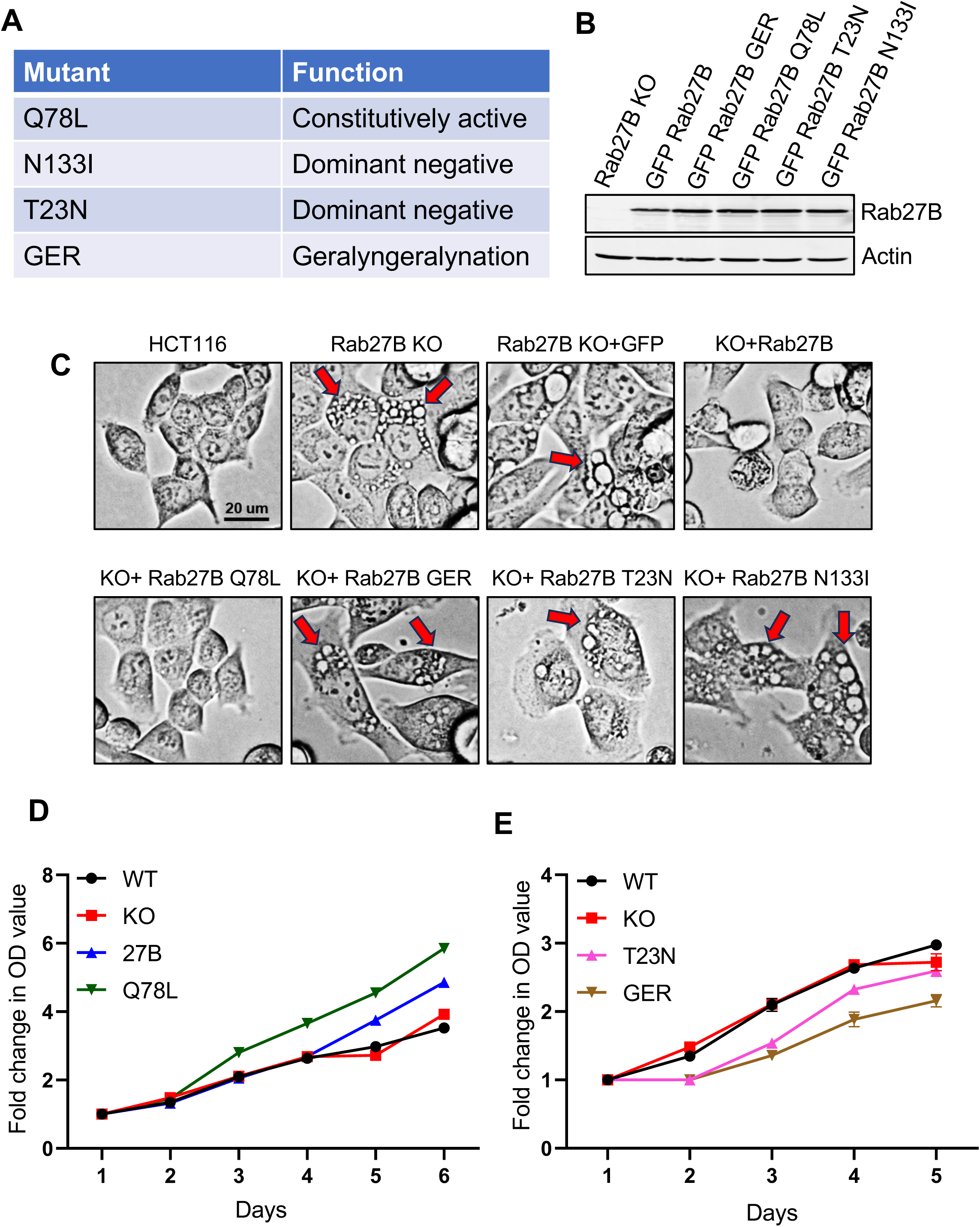
The GTPase function of Rab27B is required for autophagy role. (A, B) Rab27B KO cells were transfected with EGFP-Rab27B fusion constructs (Constitutively active Q78L mutant, dominant negative N133I and T23N mutants, and Geralyngeralynation mutant GER) as well as wt Rab27B **(A)**. Stable cell lines expressing the mutant proteins were established and confirmed by WB analysis for Rab27B **(B)**. **(C)** Brightfield images to observe the morphology and autophagic vesicle accumulation in mutant Rab27B cell lines show only wt and constitutively active (Q78L) mutant Rab27B reversed autophagy dysregulation. The dominant negative (N133I and T23N) or GER mutants failed to restore functional autophagy. **(D, E)** MTT assays were performed to assess the cell viability of mutant Rab27B cell lines over time. HCT116 wt and Rab27B KO cells were used as controls. Cells expressing constitutively active (Q78L) mutant Rab27B showed a higher proliferation rate than WT or KO cells **(D),** whereas no change was observed in the dominant negative (T23N) mutant cell line **(E)**. Three independent assays were performed.

### Effects of Rab27B depletion on CRC cell growth

Autophagy in cancer has been shown to promote tumor growth and cancer cell survival, and inhibiting autophagy can attenuate tumor cell growth. Our results show that silencing Rab27B leads to autophagic dysregulation in cancer cells, which suggests Rab27B depletion might also affect cancer cell growth. To check whether the KO cells show any growth difference, we performed MTT assays and compared the growth rate of WT and KO cells over five days. The results of MTT assay did not show any significant growth difference between the WT and KO cells (**Supplementary** Figures 4AB). Colony formation assays did not show any change in colony forming ability of the KO cells (**Supplementary** Figures 4CD).

Next we wanted to see if Rab27B deletion affects CRC cell growth *in vivo*. Recent research has shown that Rab27B can play a role in cancer stem cell growth and spheroid formation [47, 69]. So, we wanted to evaluate the growth of Rab27B KO cells in 3D culture condition. HCT116 wt and Rab27B KO cells were plated at 3-4 cells/well concentration in ultra-low adherence 96-well plates, and spheroid formation and growth was monitored over three weeks (**Figure 7A**). Interestingly, HCT116 wt cells expanded into large spheroids, while the majority of the KO cells failed to grow spheroids (**Figure 7A**). This observation was confirmed by quantification of the spheroid area using ImageJ software (**Figure 7B**). Next, we performed soft agar colony formation assays. Rab27B KO cells showed a drastic reduction in colony number (94% reduction) in the soft agar assay (**Figures 7CD**). Finally, we performed a xenograft assay with nude mice, where Rab27B KO cells failed to grow any xenograft tumors. Next, we performed a xenograft experiment to determine whether Rab27B deletion impacts tumor formation *in vivo*. 5-6 weeks old nude mice were injected with HCT116 wt and two seperate Rab27B KO clones (1 million cells/injection). Four mice per group were used with two tumors/mouse. Tumor volume was measured thrice per week and mice were harvested once the tumor volume reached 1200-1500 mm^3^. Similar to *in vitro* results, Rab27B KO cells failed to grow xenograft tumors *in vivo* (**Figure 7E**), demonstrated by the quantification of tumor growth over time (**Figure 7F**).

**Figure 7:**
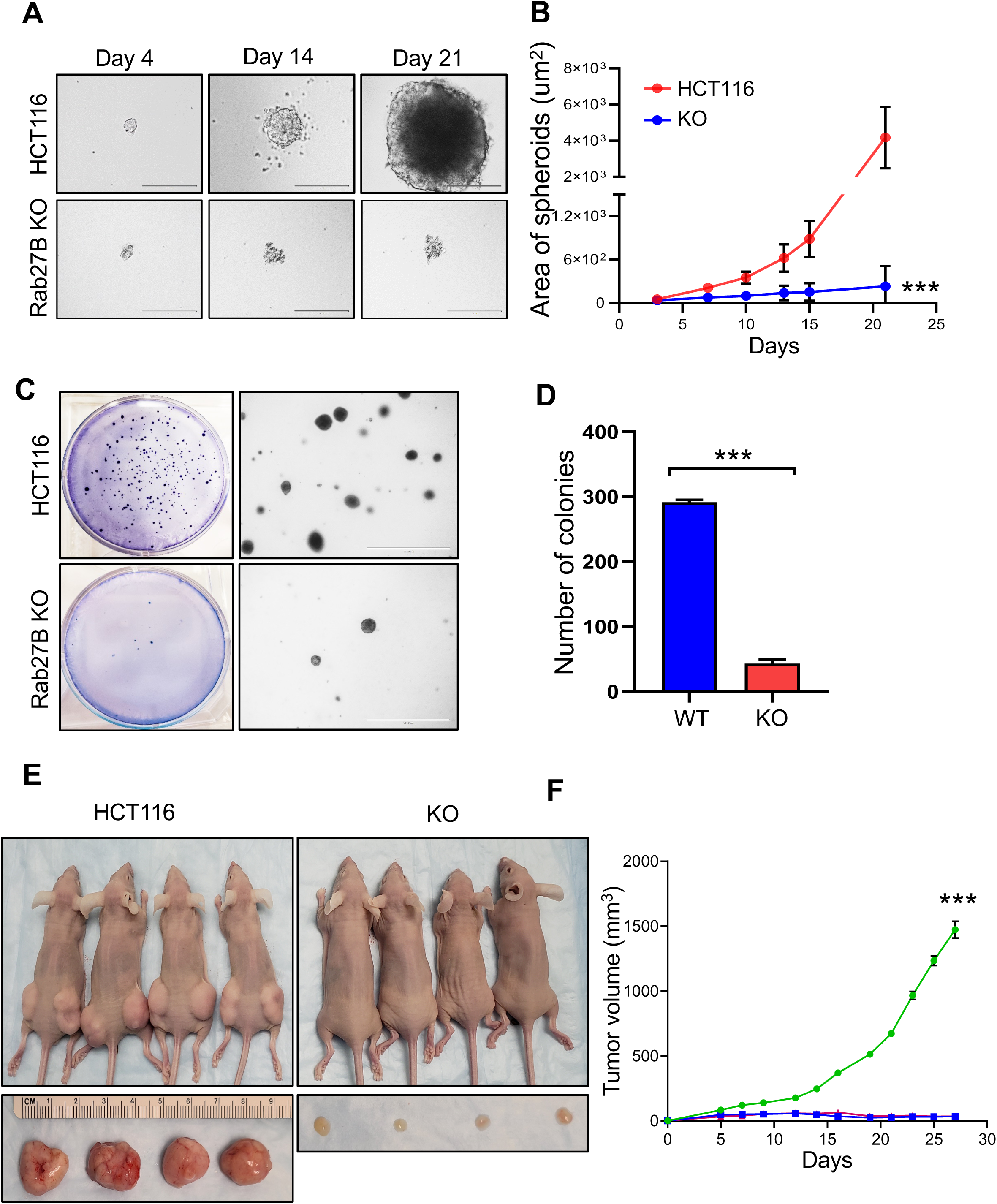
Rab27B deletion results in the inhibition of anchorage-free growth in HCT116 cells. **(A)** Brightfield images of spheroids formed by HCT116 WT and Rab27B KO cells at day 4, day 14 and day 21. **(B)**. The area of the spheroids was measured and calculated using ImageJ software. Rab27B KO cells show significant impairment in spheroid growth. **(C, D)** Rab27B deletion significantly reduces colony formation in soft agar colony formation assay. The results shown are representative of three independent experiments. **(E, F)** Rab27B deletion inhibits xenograft tumor growth *in vivo*. HCT116 wt and two separate Rab27B KO clones were injected (100 ul, 1 million cells) into the flanks of 5-6 weeks athymic nude mice (n=8 tumors /group). Tumors were measured thrice weekly. The Mice were sacrificed, and tumors were excised, weighed, measured, and photographed next to a ruler 28 days after injection. **(E)** Representative images of mice and harvested tumors from control and Rab27B KO groups. **(F)** Tumor volume were calculated by the formula- [length x (width)^2^/2]. Values given are expressed as mean ± SEM (n=8 per group). (*, *p* ≤ 0.05; **, *p* ≤ 0.01; ***, *p* ≤ 0.001).

### Rab27B KO cells are susceptible to starvation

Cancer cells deal with several stresses like scarcity of nutrition, hypoxia, immune response, etc., under *in vivo* conditions. As autophagy contributes to resistance against these stresses, we hypothesized that the autophagy-defective Rab27B KO cells should be more susceptible to these stresses. To test this hypothesis, we compared the survival percentage of HCT116 and KO cells exposed to different starvation conditions for 24 hours as follows-DMEM (4.5 g/l glucose), DMEM (1 g/l glucose), DMEM (0 g/l glucose) (**Supplementary Figure 5A**). Initial reading before and final reading after starvation were taken by MTT assay. Percent survival was calculated from the initial and the final reading. Rab27B KO cells had significantly reduced survival under stress-related serum-free conditions (**Supplementary Figure 5B**). Interestingly, we observed an increase in the Rab27B protein level following starvation in western blotting (**Supplementary Figure 5C**). This implies that Rab27B assists in tumor cell survival under conditions of stress (starvation) that are more representative of the tumor microenvironment.

Taken together, our results suggest that Rab27B is required for proper autophagy function in CRC cells. The absence of Rab27B results in a block in autophagy flux and an accumulation of autophagosomes. This dysregulation, in turn, causes suppression of tumor growth and stress response in Rab27B-depleted CRC cells (**Figure 8**).

**Figure 8:**
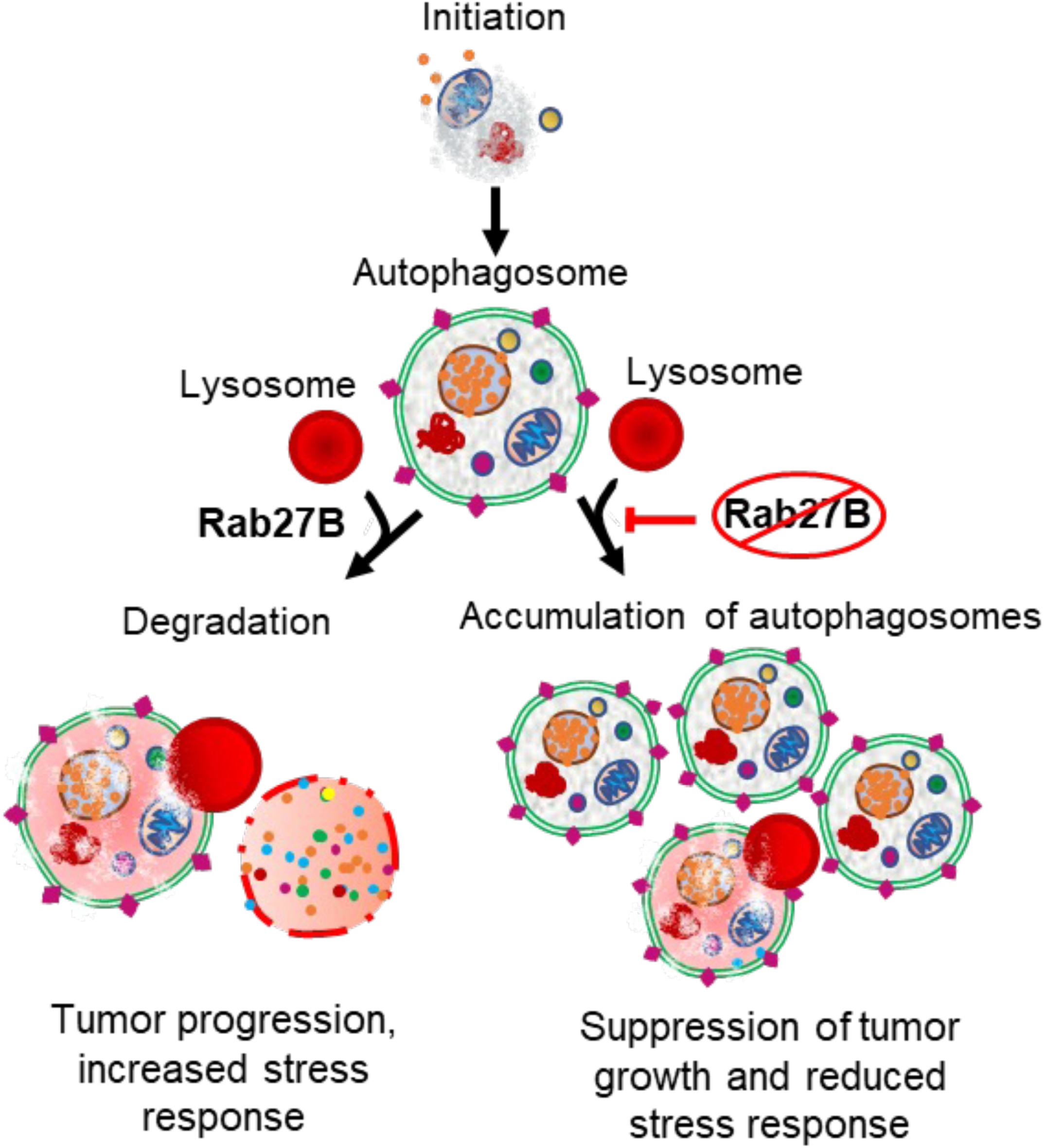
Role of Rab27B in promoting autophagy flux in CRC. Details are discussed in the text.

## DISCUSSION

In this study, we investigated a previously unstudied role of Rab27B in the regulation of autophagy in CRC. We identified that Rab27B silencing leads to a defect in autophagosome degradation, resulting in an abnormal accumulation of autophagosomes in CRC cells. Our data indicates that a block in the fusion of autophagosomes and lysosomes in the Rab27B depleted cells causes this defect in autophagy flux. This functional impact of Rab27B in the autophagy process depends on the GTP binding and geralyngeralynation of Rab27B. Furthermore, our data highlights the importance of Rab27B in the anchorage-free growth of CRC cells both *in vitro* and *in vivo*. Together, our results show the importance of Rab27B in regulating autophagy and growth of CRC cells, highlighting the therapeutic potential of Rab27B in CRC.

Rab27A and Rab27B are homolog Rabs that are major regulators of vesicle fusion and secretion in various intracellular trafficking processes. Despite having a structural similarity of 70%, Rab27A and Rab27B show diverse expression and functions in different tissues. Rab27A is overexpressed in glioblastoma, prostate, pancreas, melanoma, and thyroid cancer, whereas Rab27B shows high expression in breast, cervical, kidney, lung, colon, and stomach cancers [70]. Our previous results show that Rab27B can be overexpressed through interaction with the RNA-binding protein HuR. HuR regulates Rab27B expression post-transcriptionally by binding the 5’ UTR region of Rab27B mRNA, resulting in stabilization and subsequent protein overexpression. Here, we show that Rab27B is overexpressed in CRC tumors compared to adjacent normal tissues using a Tissue microarray containing CRC patient samples. This data is supported by previously published work that also showed that Rab27B is overexpressed in CRC and correlates with poor prognosis and survival [44, 47]. Our subsequent results show that CRC cell lines have increased expression of Rab27B to various degrees when compared with normal tissues. Interestingly, HuR does not interact with Rab27A, which agrees with the observation that Rab27A is not overexpressed in CRC.

Rab27A and Rab27B not only show diversity in their expression but also their functions. Studies revealed that, even within the same cell, Rab27A and Rab27B can perform different functions attributed to their interaction with various Rab27 effector proteins[71]. Both of these Rabs have been implicated in the proliferation, invasion, and progression of multiple cancers. One study shows Rab27B can drive cancer stem cell-like phenotype in NSCLC through extracellular vesicle secretion [69]. Their results suggest that increased EVs secreted by Rab27B overexpressing NSCLC CSCs can increase spheroid growth, clonal expansion, and invasion in bulk cancer cells. Another study revealed that Rab27B is overexpressed in CBL or JAK2 mutated myeloid malignancies and promotes cancer progression through NRAS palmitoylation and trafficking [72]. Another study showed that Rab27B promotes increased EV secretion in CRC stem cells and tumorigenicity and progression through trafficking increased mir-146a-5p through exosomes [47]. Here, we demonstrate a novel function of Rab27B in regulating the autophagy process in CRC. Rab27B silencing both by siRNA-mediated knockdown and CRISPR/cas9 knockout resulted in the accumulation of autophagosome-like vesicles in CRC cells seen both by light and Transmission electron microscopes. We confirmed that these vesicles autophagosomes by staining with autophagosome markers Cyto-ID dye and LC3 antibody, both of which showed positive staining. Rab27A depletion did not result in any autophagy defect. These results indicate that Rab27A and Rab27B may have different functions in CRC cells, excluding exosome secretion.

An accumulation of autophagosomes seen by TEM and fluorescence staining and an increase in the lipidated LC3 (LC3-II) on western blot can result from two events-a defect in autophagosome turnover, which results from a block in the later stages of autophagy or an increased autophagosome formation resulting in an inefficient turnover of autophagy flux. Our RNA-seq data did not indicate any changes in proteins involved in the initial stages of autophagy or the formation of autophagosomes. Western blot data monitoring autophagy flux did not show any change in the autophagy initiation in the Rab27B-depleted cells as well. Additionally, immunofluorescence staining with a late autophagy marker p62 revealed an accumulation of p62 positive puncta in Rab27B knockout cells. Together, these results indicate that autophagy defect in Rab27B KO cells occurs in the degradation step of the autophagosomes. This observation is in accordance with published work that shows that Rab27B KD results in accumulation of LC3-positive vesicles and indicates a possible impairment in the autophagosome-lysosome fusion [53]. One of the ways to monitor autophagy turnover is by using EGFP and mCherry-tagged LC3 protein. Our results show that Rab27B depletion in HCT116 cells expressing EGFP-mCherry-LC3 protein results in accumulation of EGFP and mCherry double-positive vesicles. In contrast, control cells with normal autophagy flux show increased mCherry-positive vesicles. Together these results indicate that there is indeed a defect in the turnover of autophagosomes when Rab27B is silenced in CRC cells. This defect in the turnover can occur at the fusion stage (autophagosome-lysosome fusion) or inefficient cargo degradation once fusion has occurred. Our results from the lysotracker experiment indicate that autophagic vesicles in Rab27B KO cells are not lysotracker positive in response to induction of autophagy, whereas control cells showed lysotracker positive autophagic vesicles. Together these results show that Rab27B depletion impairs autophagosome-lysosome fusion, implicating that Rab27B functions in the fusion stage of autophagy.

Rab27B functions through GTPase activity and switches between active GTP bound and inactive GDP bound conformations. Rab27B also undergoes post-translational modification by geralyngeralynation, which acts as an anchor for specific membrane binding interactions. Even though Rab proteins are well studied for their GTPase activity, research has shown that Rab can function through other parts of their structures, too. Although Rab27B has been functionally implicated in a number of different pathways, the functional necessity of its GTPase activity has not been studied extensively except for one study that reported that Rab27B promotes the growth, invasion, and metastasis in ER-positive breast cancer in GTP-binding and geranyl-geranyl dependent manner [73]. They generated mutant forms of Rab27B that encoded Rab27B-T23N and Rab27B-N133I, which are defective in GTP binding; thus, inactive, Rab27B-Q78L, which is defective in GTP hydrolysis, therefore constitutively active and Rab27B-GER which is a geranyl-geranyl binding mutant. Here, in HCT116 Rab27B KO background, we stably expressed these mutant forms of Rab27B and saw a rescue of the autophagy defect only in the WT and constitutively active Q78L mutant Rab27B expressing cells. GTP binding mutants T23N and N133I, as well as geralyngeralyn mutant GER, did not show any recovery of the autophagy dysregulation. Overall, these results indicate that the autophagy function of Rab27B is dependent on its GTPase activity and geranylgeranyl modification.

Interestingly, whereas Rab27B deletion does not affect the proliferation or colony formation ability of CRC cells *in vitro* in 2D culture, we do see a striking growth deficiency *in vitro* 3D culture as well as *in vivo* xenograft study. In cell culture, Rab27B KO cells failed to grow spheroids and showed a 94% reduction in soft agar colony formation assay. These results highlight the importance of Rab27B in anchorage-independent growth, which is characteristic of cancer stem cells. This is in line with other studies that show that Rab27B plays a key role in CSC growth and proliferation. One study showed that Rab27B is overexpressed, and depletion of Rab27B resulted in growth reduction of NSCLC stem-like cells but did not have any growth effects on the bulk cancer cells [69]. Research shows that cancer stem cells require high autophagy flux, so defective autophagy could drastically affect these cells. We show that nude mice injected with Rab27B KO cells grew drastically smaller tumors than those injected with HCT116 wt cells. This result is coherent with another study that showed that HCT116 cells do not show growth defects in response to autophagy impairment *in vitro* but show a significant reduction in xenograft tumor growth *in vivo* [74]. This growth phenotype most likely indicates a higher requirement of autophagy *in vivo* condition. Even though Rab27B KO cells do not show growth reduction *in vitro*, Rab27B KO cells stably expressing Rab27B-Q78L mutant showed a significant increase in proliferation compared to parental Rab27B KO cells or HCT116 wt cells.

Overall, our results elucidate a mechanism by which Rab27B promotes anchorage-free cell and tumor growth in CRC by regulating autophagy. While the outcome of our study indicates that Rab27B is involved in the autophagic process and promotes autophagic flux, the exact mechanism of how Rab27B plays this role remains to be explored. This mechanism can be explored in future studies by looking at effector proteins involved in Rab27B function, as well as the binding of Rab27B to key protein complexes involved in the autophagy process. So far, eleven Rab27-specific effectors have been identified. We predict that Ra27B interacts with different effector proteins to regulate the autophagy process or exosome secretion. Our study also underscores the promising therapeutic potential of Rab27B through autophagy inhibition. Future investigations will focus on discovering specific inhibitors against Rab27B for CRC treatment.

## SUPPLEMENTARY FIGURES

**Supplementary Figure 1:**
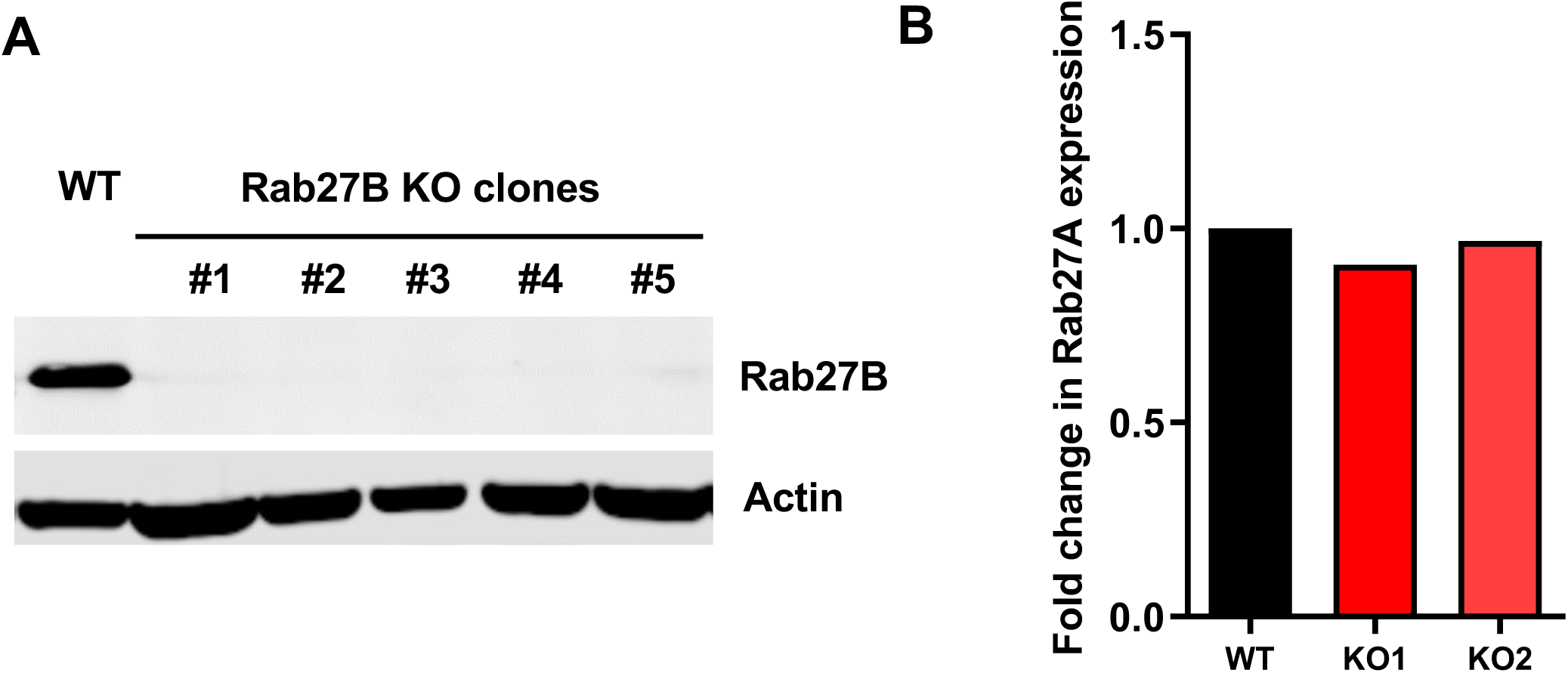
Rab27B knockout HCT116 cells. **(A)** Individual Rab27B KO clones were isolated and screened by WB for the loss of Rab27B protein. Actin was used as loading control. **(B)** RT-PCR was performed to measure Rab27A expression in Rab27B KO cells.

**Supplementary Figure 2:**
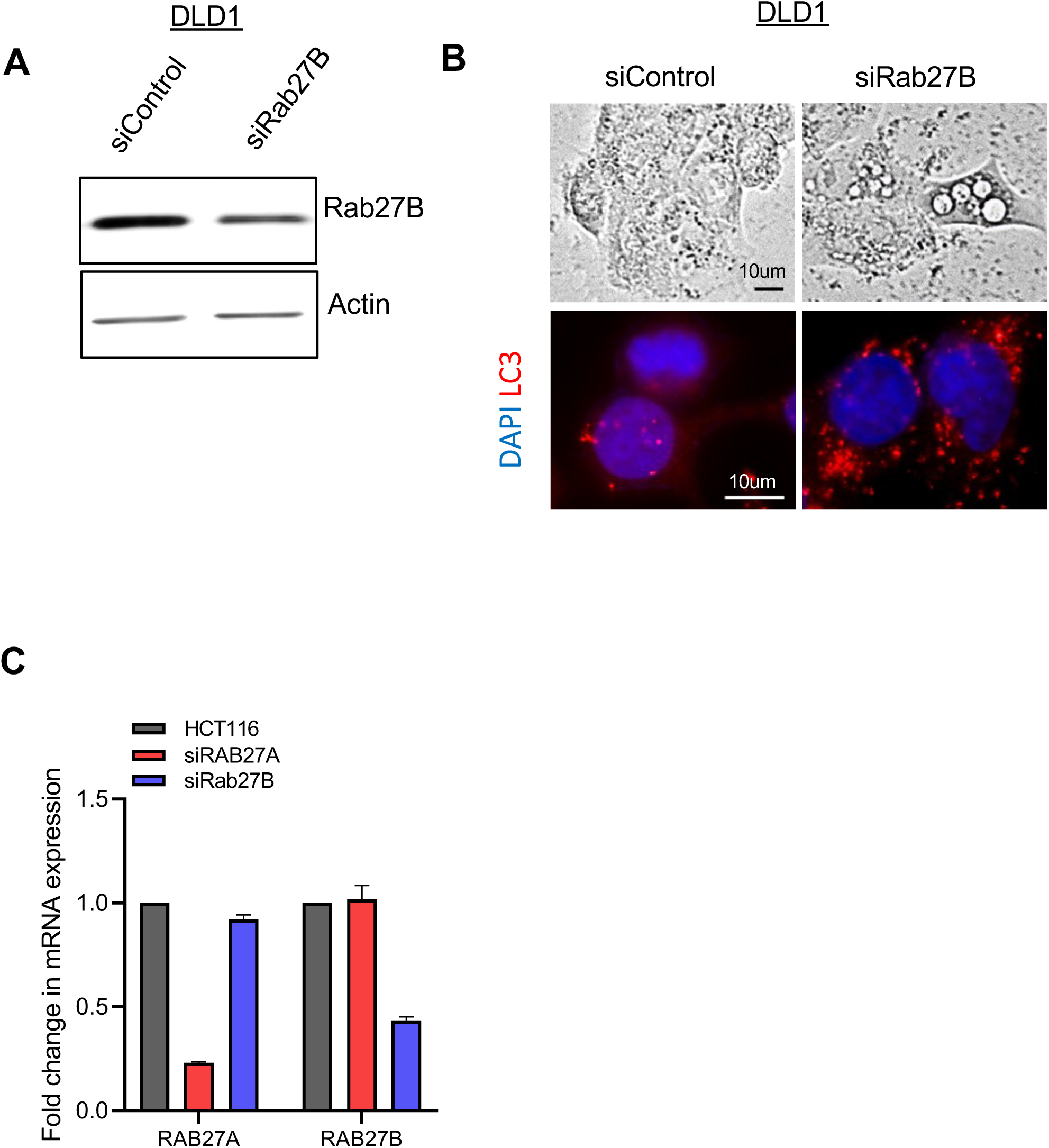
LC3 immunostaining of Rab27B knockdown DLD1 cells. **(A)** WB validation of Rab27B Knockdown in DLD1 cells. **(B)** Rab27B depletion resulted in LC3 positive vesicles (red) in DLD1 cells. **(C)** Rab27A and Rab27B knockdown was validated by RT-PCR. The data shown are representative of three independent experiments. (*, *p* ≤ 0.05; **, *p* ≤ 0.01; ***, *p* ≤ 0.001).

**Supplementary Figure 3:**
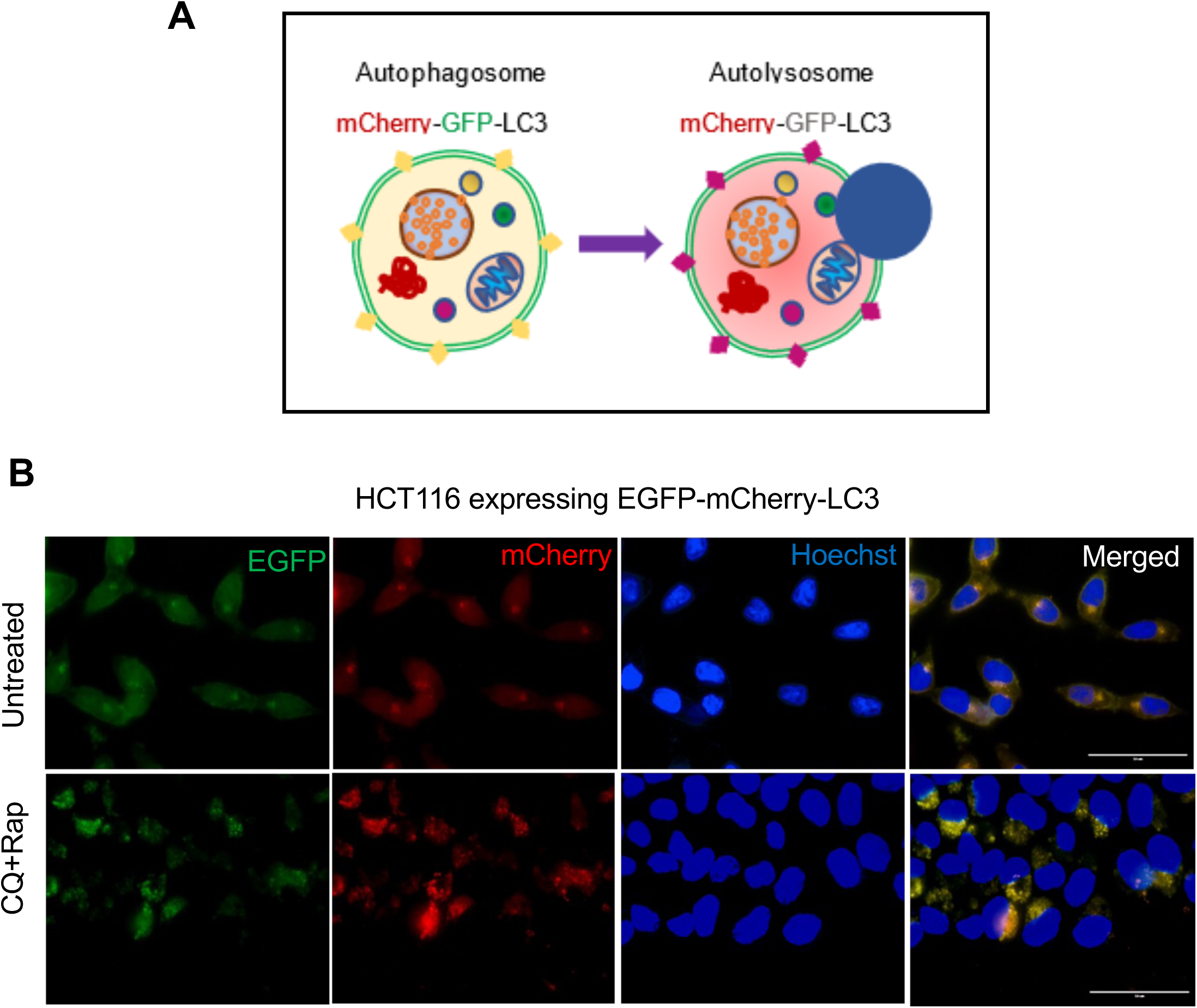
mCherry-GFP-LC3 in the assessment of autophagy flux. **(A)** Following transfection with mCherry-GFP-LC3 reporter plasmid, mCherry-GFP-LC3 localizes to autophagosome membranes. So, autophagosomes are double positive for mCherry and GFP (indicated by yellow puncta when red and green channels are merged). Autolysosomes are positive only for mCherry due to the degradation of GFP following lysosomal fusion (indicated by red puncta). **(B)** HCT116 stably expressing EGFP-mCherry -LC3 fusion protein was created. Untreated control cells show mostly cytosolic LC3 (yellow). Cells treated with rapamycin (0.5 um) and chloroquine (10 um) show the formation of yellow and red autophagic vesicles.

**Supplementary Figure 4:**
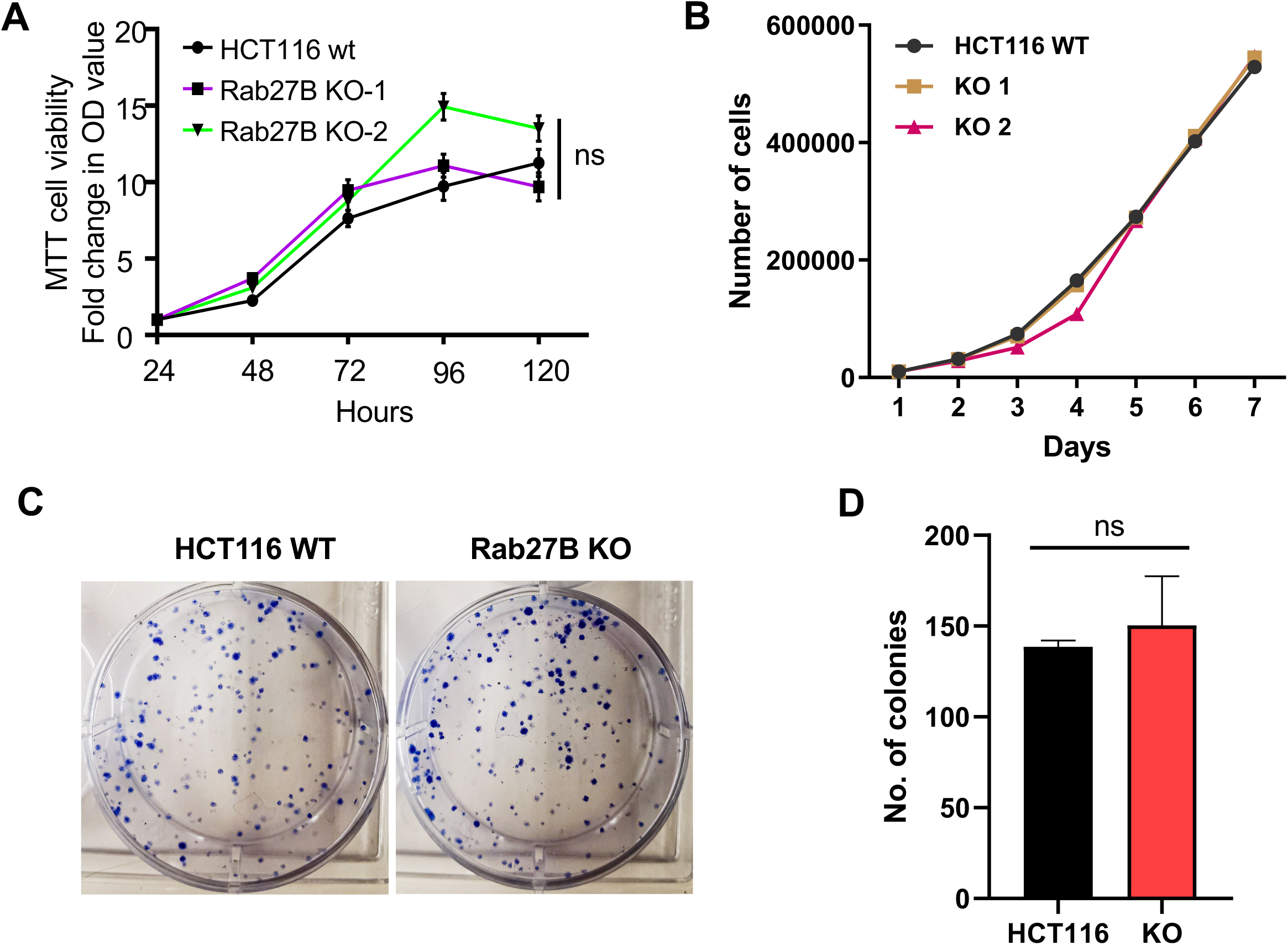
Rab27B KO cells do not show any change in proliferation in 2D cell culture. **(A)** Cell proliferation of HCT116 and Rab27B KO cells compared in normal growth media (DMEM with 10% FBS). Cell proliferation was measured by MTT assay normalized to the initial OD value. **(B)** HCT116 and Rab27B KO cells were seeded in 6 well plates, and cell counting was done every day for seven days using KOVA Glasstic Slides. **(C)** Colony formation assays were performed to compare the colony-forming ability of HCT116 and Rab27B KO cells. **(D)** The number of colonies was counted under the microscope. Experiments were done in triplicates and repeated three times. The result is shown as mean ± SEM. (*, *p* ≤ 0.05; **, *p* ≤ 0.01; ***, *p* ≤ 0.001).

**Supplementary Figure 5:**
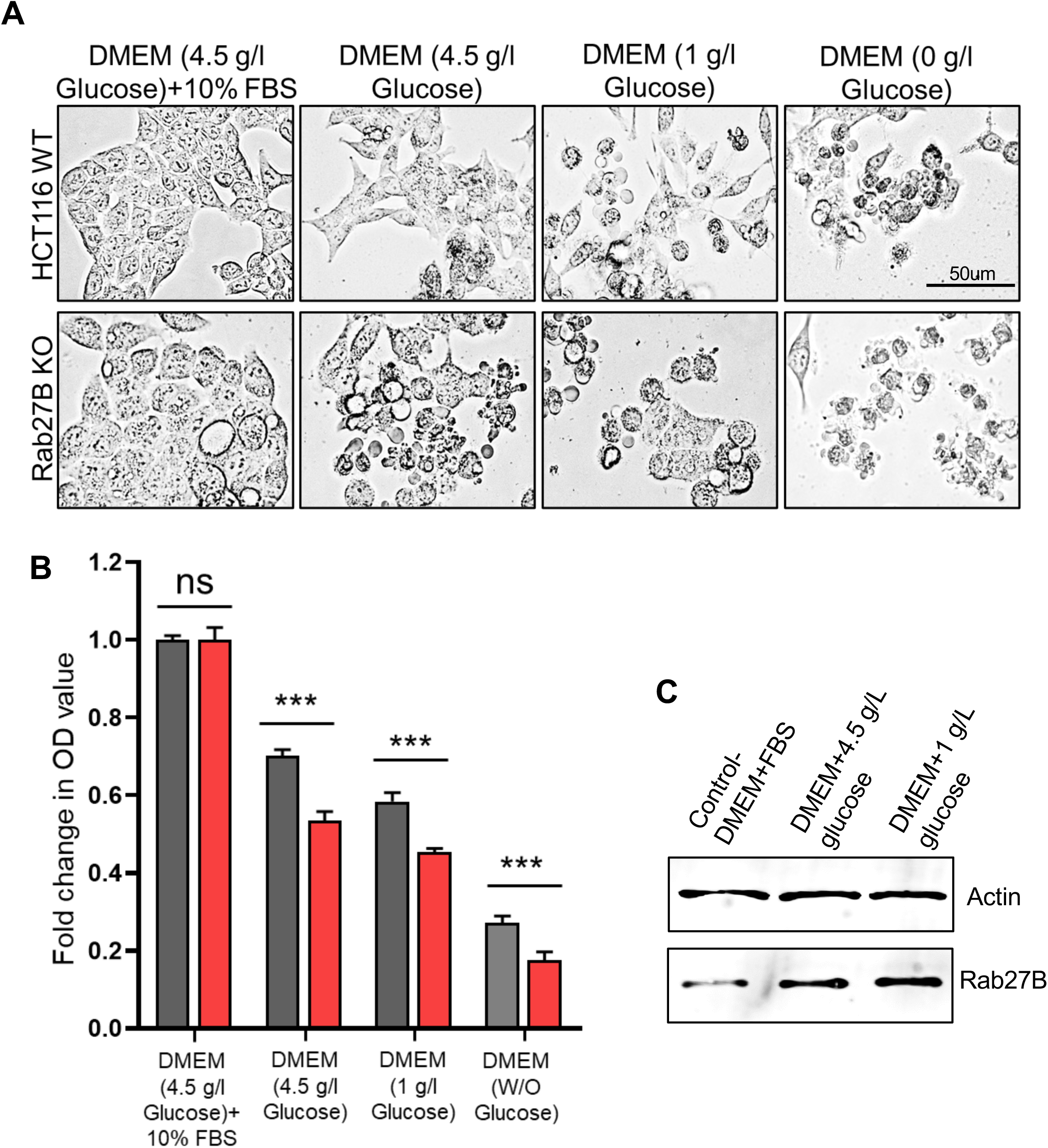
Rab27B KO cells show reduced survival percentage following starvation. **(A)** HCT116 and KO cells were grown in 96-well plates containing DMEM with 10% FBS for 48 hours, followed by serum starvation in only DMEM (4.5 g/l glucose), DMEM with low glucose (1g/l glucose) and DMEM without any glucose for 24 hours. (B) Cell survival was measured by MTT assay normalized to the initial OD value before starvation. (C) Western blotting shows an increase in Rab27B protein level following starvation. Actin was used as the loading control. Experiments were repeated three times. The results are shown as mean ± SEM. (*, *p* ≤ 0.05; **, *p* ≤ 0.01; ***, *p* ≤ 0.001).

## Notes

### Competing Interest Statement

The authors have declared no competing interest.

